# A Statistical Framework to Predict Functional Non-Coding Regions in the Human Genome Through Integrated Analysis of Annotation Data

**DOI:** 10.1101/018093

**Authors:** Qiongshi Lu, Yiming Hu, Jiehuan Sun, Yuwei Cheng, Kei-Hoi Cheung, Hongyu Zhao

## Abstract

Identifying functional regions in the human genome is a major goal in human genetics. Great efforts have been made to functionally annotate the human genome either through computational predictions, such as genomic conservation, or high-throughput experiments, such as the ENCODE project. These efforts have resulted in a rich collection of functional annotation data of diverse types that need to be jointly analyzed for integrated interpretation and annotation. Here we present GenoCanyon, a whole-genome annotation method that performs unsupervised statistical learning using 22 computational and experimental annotations thereby inferring the functional potential of each position in the human genome. With GenoCanyon, we are able to predict many of the known functional regions. The ability of predicting functional regions as well as its generalizable statistical framework makes GenoCanyon a unique and powerful tool for whole-genome annotation. The GenoCanyon web server is available at http://genocanyon.med.yale.edu

## Introduction

Annotating functional elements in the human genome is a major goal in human genetics. Despite years of efforts from both experimental and computational scientists, functional annotation remains challenging, especially in the non-protein-coding regions. It is estimated that approximately 98% of the human genome is non-protein-coding^1^. Because of the apparent importance of coding regions, many computational tools have been developed to annotate DNA variants in the coding regions^2-4^. Although the non-coding regions were considered “junk DNA” for many years, much has been learned on the potential roles of these regions in the last decade. First, extensive comparative genomic studies have shown that the majority of mammalian-conserved regions consist of non-coding elements^5^. Second, results from genome-wide association studies show that close to 90% of the significant variants associated with human diseases reside outside of the coding regions^6^, only slightly less underrepresented among all the variants in the human genome, where about 95% of known variants are from the non-coding regions. Third, high-throughput experiments, e.g. the ENCODE project^7^, also suggest that a large fraction of the human genome are functionally relevant. All of this evidence suggests the importance and need for extending the annotation tools from the coding regions to the entire human genome.

Despite the increasing need to functionally annotate the human genome, there is no universal definition of genomic function^8,9^, which differs among geneticists, evolutionary biologists, and molecular biologists. The experimental approaches and analysis techniques of detecting functional genomic elements among these scientists also vary greatly. Extensive work in some genomic regions such as the β-globin gene complex has shown that no single approach is sufficient to identify all the regulatory activities in the non-coding regions^8,10^. In order to obtain a comprehensive picture of the genomic functional structure, all the valuable information acquired through different approaches needs to be combined using appropriate statistical learning techniques.

Several annotation tools focusing on the non-coding regions have been established recently^11-15^. Similar to the long list of deleteriousness prediction tools developed for the coding regions, most of these new methods aim to distinguish tolerable variants from the deleterious ones. Though important, prediction of deleteriousness does not cover every aspect of functional annotation. The potential of these variant classifiers in understanding the genomic architecture on a large scale and in detecting regulatory elements such as cis-regulatory modules remains to be thoroughly investigated. Moreover, scientists now routinely analyze different cell types^7^, and even single cells^16^. In order to keep up with these technological advances, it is critical to develop a functional annotation framework that can be generalized to different species, cell types, and single cells. Such a generalizable framework can be achieved through biologically-motivated and statistically-justified models. As for choosing between a supervised approach, where some gold standard datasets are needed to train the model, and an unsupervised approach, where no labeled data are used, we focus on developing an unsupervised learning method in this article. This is because current supervised-learning-based annotation tools suffer from highly biased training data, which is largely due to our limited knowledge of non-coding regions. This may become less of an issue after we have gained a deeper understanding of non-coding functional mechanisms. However, at such an early stage, we think unsupervised learning techniques would be advantageous.

In this paper, we present GenoCanyon (inspired by the canyon-like plots it generates), a whole-genome annotation tool based on unsupervised statistical learning. From a collection of the comparative genomic conservation scores and biochemical signals obtained from the ENCODE project^17^, the posterior probability of a genomic position being functional is used as the prediction score. Compared to existing methods, GenoCanyon not only measures the deleteriousness of variants, but also the functional potential of each genomic location. Its flexible and generalizable statistical framework could also benefit future applications.

## Results

### Estimating the Proportion of Functional Regions in the Human Genome

Genetic approaches that focus on studying the consequences of genetic perturbations are often referred to as a gold standard for defining function^8^. Such a genetic definition is also directly related to causal inference, which is at the core of developmental biology and disease research^9^. In this study, we also adopt this genetically meaningful definition of genomic function. On the other hand, we treat the conservation measures and the biochemical signals as consequences of genomic function (Figure 1A). For a specific location in the human genome, define Z to be the latent indicator of function. We collected 22 different annotations, denoted as ***A*** (Supplementary Table 1). We also assumed that the 22 annotations are conditionally independent given Z (Figure 1B). Then, the posterior probability (*P Z* = 1 ***A***) serves as the prediction score of the functional potential at this location (See **Methods**, Figure 1C).

**Figure 1.**
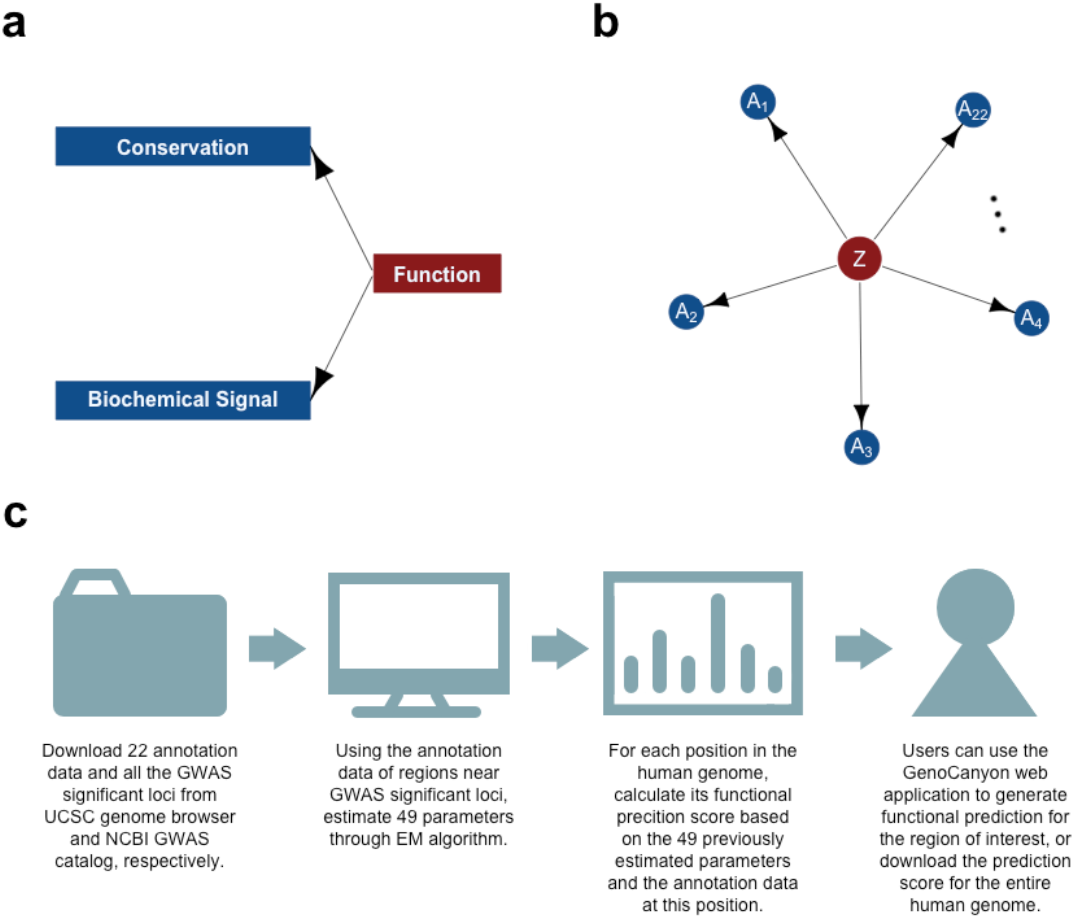
Modeling of causal relationship among variables. **a)** We adopt the biologically meaningful definition of function, and treat conservation measures and biochemical signals as consequences. **b)** The latent functional indicator Z is modeled as the parental variable and all the 22 annotations are treated as consequences. Also, we assume there is no direct causal relationship between any two annotations. Therefore the annotations are conditionally independent given Z. **c)** Workflow of GenoCanyon functional prediction.

We have pre-calculated the prediction scores for the entire human genome (hg19). Overall, when using 0.5 as the cutoff for defining functionality, 33.3% of the human genome was predicted to be functional. The proportion of functional elements is mostly stable across chromosomes (Supplementary Table 2; Supplementary Figure 1). We note that the functional proportion of the human genome has been estimated using many different approaches^8,18-22^ and results differed drastically. Comparative genomic analysis of multiple mammals revealed that constrained elements consist of approximately 4.5% of the human genome^18,19^. At the other extreme, the ENCODE project found that 80% of the human genome has detectable biochemical activities in at least one cell line^7^.

However, it has been discussed recently that several corrected constraint estimations would each suggest two to three times increase to the original estimate of 4.5%^20-22^. Also, it still remains non-trivial to distinguish real biochemical signals from biological noises in the ENCODE data^8^. The large amount of observed biochemical activities have also been criticized to be more like an “effect” rather than “function”^9^. Our prediction falls in the middle of these highly diverging estimates of functional regions in the literature. It is worth noting that the GenoCanyon functional prediction represents a mixed probability involving multiple tissues. A smaller proportion of the human genome would be expected to be functional for a particular tissue.

### Prediction for cis-regulatory Modules in the HBB Gene Complex

The intensively studied β-globin (HBB) gene complex on chromosome 11 contains embryonically expressed HBE1, fetally expressed HBG1 and HBG2, and adult globin genes HBD and HBB, along with a pseudogene HBBP1. This locus is known to provide a paradigm for developmental gene expression and regulation^23,24^. A large number of cis-regulatory modules (CRMs) that control both the developmental timing and the spatial pattern of gene expression have been discovered in the HBB gene complex^10^. More interestingly, the epigenetic and evolutionary signals at these CRMs differ substantially^8^. Therefore, the HBB gene complex provides a perfect example to test if GenoCanyon could effectively combine different sources of signals and successfully predict the functional segments.

We analyzed the prediction results in the HBB gene complex. On the entire chromosome 11, 32.2% of the DNAs were predicted to be functional. Strong enrichment of signals was observed at this locus. Using 0.5 as the cutoff, 62.2% of HBB gene complex and 97.0% of the CRMs were predicted as functional (Figure 2A). Remarkably, a cluster of five DNase I hypersensitive CRMs upstream of the HBB gene complex, known as the locus control region (LCR)^25^, showed strong functional signals as a whole (Figure 2B). The 3’HS1 enhancer blocker (chr11: 5226013-5226493; hg19) downstream of the HBB gene complex was also successfully predicted with high resolution. Interestingly, these CRMs showed highly variable patterns of annotations (Figure 2C). This proved that GenoCanyon could effectively combine different sources of information. Recent research revealed several new regulatory elements at this locus, including one in the intergenic region between HBBP1 and HBG1^24^, and another one upstream of HBD^23^. These elements also reside in the highly scored regions. Moreover, it is worth noting that the understanding of CRMs is still incomplete even in a relatively well-studied region such as the HBB complex. Some of the apparent false positives might actually be regulatory elements not yet discovered. The functional regions provided by our method could potentially offer a guideline for further studies.

**Figure 2.**
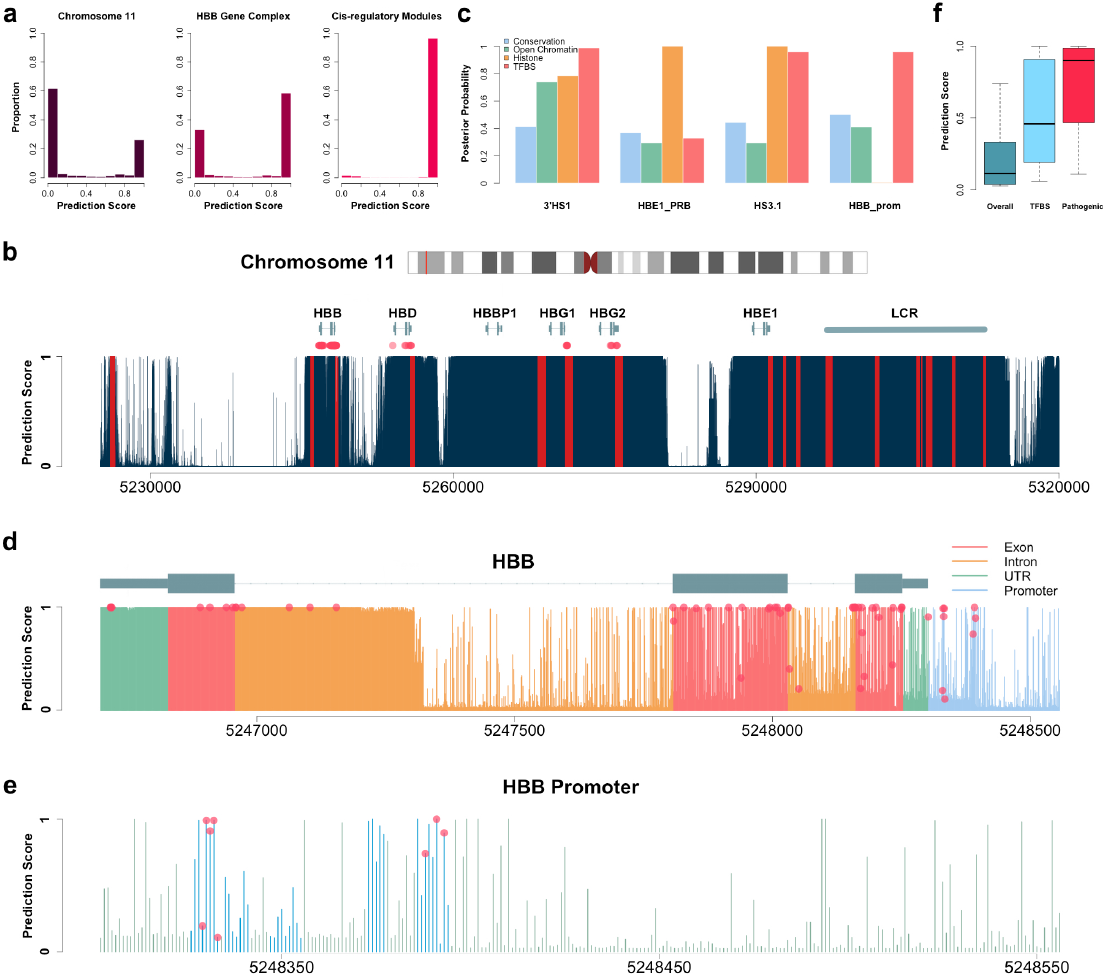
Functional prediction for the HBB gene complex. **a)** Histogram of the prediction scores in chromosome 11, HBB gene complex, and the 23 CRMs. 32.2%, 62.2% and 97.0% are predicted as functional, respectively. **b)** Prediction results for the HBB complex. Dark blue bars show the prediction score at each location. All the 23 CRMs are marked in red. There appears to be fewer than 23 red bars because some of the CRMs are very close to each other. Red dots indicate the locations of known pathogenic SNPs downloaded from the NCBI Variation Viewer. **c)** The posterior probabilities given a single group of annotations could be used to measure the relative contribution of different sources of information (See **Methods**). Four CRMs are plotted to illustrate that prediction scores are driven by different annotations in different CRMs. **d)** Prediction results for the HBB gene and its promoter. The promoter, UTRs, introns and exons are marked with different colors. Red dots show the prediction scores of the pathogenic variants. **e)** Prediction results for the HBB promoter. Known protein binding sites in the HBB promoter are marked in blue. Red dots show the prediction scores of the pathogenic variants. **f)** Boxplot of the prediction scores of HBB promoter, known protein binding sites, and pathogenic variants.

Among the 23 CRMs being reviewed^10^, only the promoter of HBB did not get the perfect score (Table 1). Therefore, we analyzed the HBB gene and its promoter in more details (Figure 2D). Within the HBB gene, the 600bp segment near the 3’UTR was predicted to be functional. 77 pathogenic or likely pathogenic SNPs were downloaded from the NCBI Variation Viewer (http://www.ncbi.nlm.nih.gov/variation/view/). Interestingly, 14 of these pathogenic SNPs, including 4 in the 3’UTR and 6 in the second intron, lie in this 600bp functional segment. In the upstream half of the second intron, prediction scores were substantially lower. No pathogenic variants could be found in that region. Overall, using 0.5 as the cutoff, 89.6% (69 out of 77) of the pathogenic SNPs located at functional locations. Within the HBB promoter, 75% (6 out of 8) of the pathogenic variants located at functional locations (Figure 2E). Moreover, bumps of high scores could be observed at the known protein binding sites in the HBB promoter^26^. When comparing the entire HBB promoter, known protein binding sites, and the pathogenic variants within the promoter, there was a substantial increase in prediction score (Figure 2F). All of this evidence suggests that important functional segments could still be detected locally even in a generally lower-scored region.

**Table 1.** Mean prediction scores of the known CRMs in the HBB gene complex.

| **Name** | **Start** | **Stop** | **Mean Score** |
| --- | --- | --- | --- |
| HS5 | 5312534 | 5312694 | 1.000 |
| HS4 | 5309419 | 5309707 | 1.000 |
| HS3.2 | 5306814 | 5307392 | 1.000 |
| HS3.1 | 5306356 | 5306418 | 1.000 |
| HS3 | 5305882 | 5306169 | 1.000 |
| HS2_neg | 5302090 | 5302174 | 1.000 |
| HS2_pos | 5301795 | 5302089 | 1.000 |
| HS1 | 5296894 | 5297517 | 1.000 |
| HBE1_NRA | 5294082 | 5294308 | 1.000 |
| HBE1_PRA | 5293982 | 5294081 | 1.000 |
| HBE1_NRB | 5292886 | 5292928 | 1.000 |
| HBE1_PRB | 5292690 | 5292886 | 0.999 |
| HBE1_up | 5291344 | 5291610 | 1.000 |
| HBE1_prom | 5291175 | 5291343 | 1.000 |
| HBG2_up | 5276215 | 5276745 | 1.000 |
| HBG2_prom | 5276011 | 5276214 | 1.000 |
| HBG1_up | 5271291 | 5271813 | 0.999 |
| HBG1_prom | 5271086 | 5271290 | 1.000 |
| HBG1_3'enh | 5268365 | 5269114 | 1.000 |
| HBD_prom | 5255713 | 5256160 | 1.000 |
| HBB_prom | 5248301 | 5248556 | 0.253 |
| HBB_3'enh | 5245876 | 5246140 | 1.000 |
| 3'HS1 | 5226013 | 5226493 | 0.987 |

\*Coordinates are based on hg19.

### Prediction for ZRS, an enhancer of the SHH gene

Zone of polarizing activity regulatory sequence (ZRS) is one of the most studied developmental enhancers. It is located in the fifth intron of the protein-coding gene LMBR1, approximately 1Mb upstream of SHH’s transcriptional start site^27,28^. Through linkage mapping of several large families with preaxial polydactyly (PPD) and triphalangeal thumb, an associated locus of approximately 500 kb was identified on chromosome 7q36. In later studies, the region was further narrowed down to the fifth intron of LMBR1^29-31^. As a highly conserved 774 bp region in this intron, ZRS has been intensively studied. It has been shown to be crucial for limb development not only in humans, but also in mice, dogs, cats, and even chickens^27^.

We investigated the prediction results in gene LMBR1. A highly scored plateau could be observed in its fifth intron (Figure 3A). The mean predicted score for this intron was 0.595. This was higher than the mean score of the entire LMBR1 transcript (0.385), of all the introns in LMBR1 (0.384), and even of all the exons in LMBR1 (0.448). These results showed strong signs of function in the fifth intron. The ZRS region got an even higher mean predicted score 0.871, which confirmed its importance (Figure 3B). When observing its surrounding region, ZRS could be easily identified as a dense region with high prediction scores (Figure 3C). Moreover, the ZRS region serves as one of the most well studied examples for pathogenic variants in an enhancer. A total of 13 single nucleotide variants in ZRS have been identified to cause human limb malformations^27^. All these 13 SNVs were predicted to be highly functional, with the mean prediction score 0.987 (Figure 3D).

**Figure 3.**
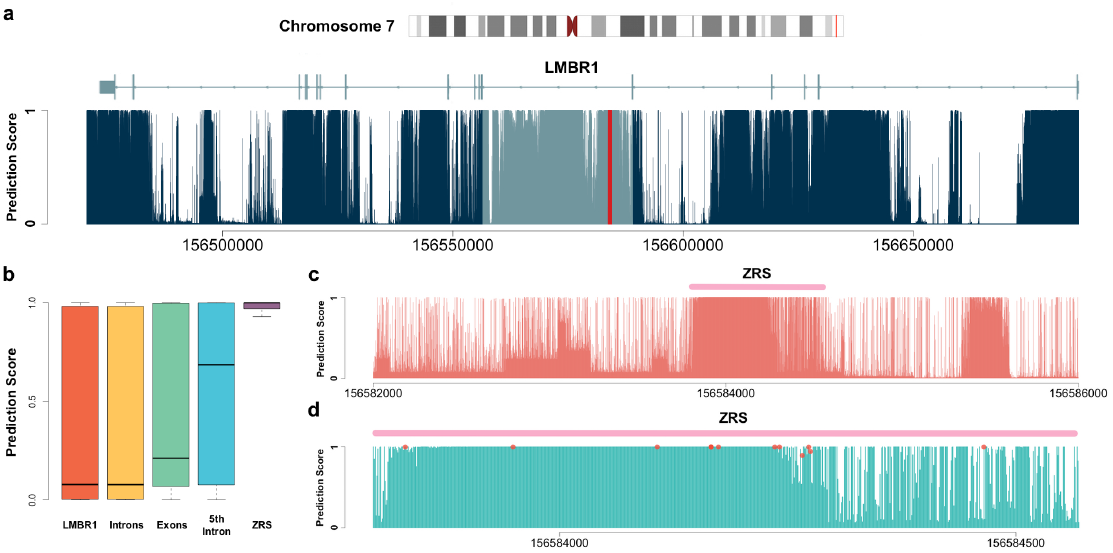
Prediction results for the SHH enhancer in LMBR1. **a)** Prediction scores in the LMBR1 gene. The fifth intron and ZRS are highlighted in light blue and red, respectively. **b)** Boxplot of the prediction scores in LMBR1, 16 introns, 17 exons, the 5th intron, and ZRS. The results highlighted the function in the 5th intron of LMBR1 and confirmed the importance of ZRS. **c)** Prediction results for the surrounding region of ZRS, which is highlighted in pink. An obvious highly scored plateau can be observed at ZRS. **d)** The prediction results within the ZRS. 13 pathogenic variants are discovered in ZRS. The predicted scores at their locations are marked with red dots. There appears to be only 11 dots because three variants all reside at location 156584166 (hg19).

In conclusion, our method successfully identified the fifth intron of LMBR1 as a functional region. It also further confirmed the importance of ZRS. It is notable that the large number of identified pathogenic variants in ZRS is possibly subject to the ascertainment bias. In fact, mutations in ZRS did not account for the limb malformation in all the studied families^32^. Our prediction in the surrounding regions has the potential to guide future studies.

### Prediction for Functional Elements in the Human X-inactivation Center

X-chromosome inactivation, originally described 50 years ago^33^, is the mechanism for X-chromosome dosage compensation in mammals. The long non-coding RNA Xist has been shown to be both necessary and sufficient to induce X-chromosome inactivation in mouse ES cells^34^. The surrounding genomic region, often referred to as the X-inactivation center (Xic for mouse and XIC for human), contains several crucial regulatory elements for mouse X-inactivation^35^. However, recent studies have suggested the existence of substantial variations in the mechanism of achieving X-inactivation among species^36-39^. We applied GenoCanyon on human XIC to predict the functional potential of the orthologs of known regulatory elements in mouse models (Figure 4A; Table 2).

**Figure 4.**
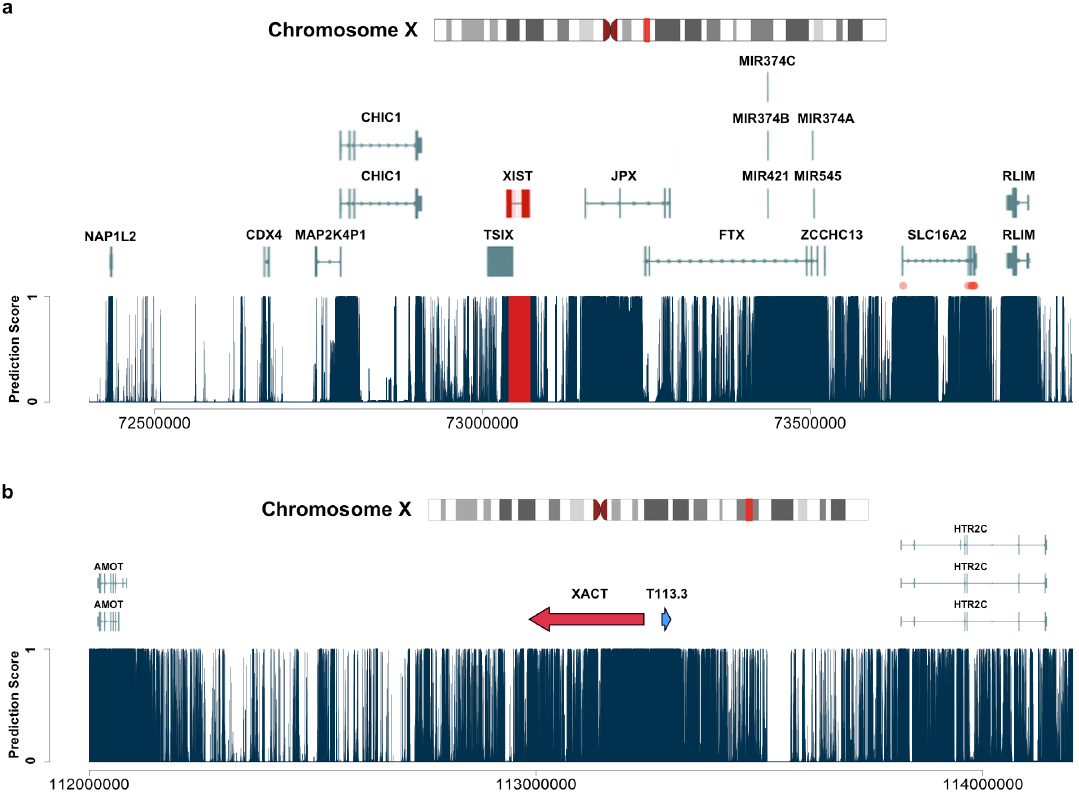
Prediction results for regions involved in human X-inactivation. Each dark blue line shows the prediction score at a single base. **a)** Functional prediction for the human XIC. All the RefSeq transcripts in this region are plotted. The master lncRNA XIST is highlighted in red. Red dots show the locations of known pathogenic variants downloaded from the NCBI variation viewer. **b)** Functional prediction for the intergenic region between AMOT and HTR2C on chromosome Xq23. A red and a blue arrow represent the recently discovered transcripts XACT and T113.3, respectively.

**Table 2.** Prediction results for the 6 protein-coding genes, 4 lncRNA genes, and 1 pseudogene in the human XIC, as well as all their transcripts in RefSeq.

| Gene Name | Gene Mean Score | Transcript ID | Transcript Mean Score |
| --- | --- | --- | --- |
| NAP1L2 | 0.988 | NM_021963.3 | 0.988 |
| CDX4 | 0.234 | NM_005193.1 | 0.554 |
| CHIC1 | 0.229 | NM_001039840.2 | 0.931 |
|  |  | NM_001300884.1 | 0.955 |
| ZCCHC13 | 0.184 | NM_203303.2 | 0.184 |
| SLC16A2/XPCT | 0.575 | NM_006517.4 | 0.951 |
| RLIM/RNF12 | 0.973 | NM_016120.3 | 0.930 |
|  |  | NM_183353.2 | 0.930 |
| XIST | 0.999 | NR_001564.2 | 0.998 |
| TSIX | 0.383 | NR_003255.2 | 0.383 |
| JPX | 0.501 | NR_024582.1 | 0.256 |
| FTX | 0.438 | NR_028379.1 | 0.304 |
| MAP2K4P1 | 0.161 | NR_029423.1 | 0.095 |

Xist and its antisense ncRNA Tsix, as well as two upstream ncRNAs Ftx and Jpx have all been shown to haven cis-regulatory roles in mouse X-inactivation^40-43^. Our prediction confirmed the function of the master ncRNA XIST in human. Both the XIST gene and its transcribed regions got nearly perfect prediction scores. Moreover, a XIST-specific peak of high score could be observed on Figure 4A, showing satisfying resolution of prediction. Studies suggesting a truncated form of TSIX in human have led to some debate in its function. Compared to its mouse ortholog, the human TSIX gene has lost the CpG island as well as the enhancer elements Dxpas34 and Xite^44^. In our prediction, TSIX got mean score 0.383, which is low for such an active genomic region. When considering only the region that does not overlap with XIST, the number even dropped to 0.197. A recently discovered lncRNA, Linx, has been hypothesized to take part in Tsix expression in mice^45^. In the mouse genome, the Linx gene lies between two protein-coding genes Nap1l2 and Cdx4. However, its human ortholog has not yet been discovered. The intergenic region between human NAP1L2 and CDX4 has a low mean prediction score 0.016, which argues against not only the existence of LINX in human, but also TSIX function. Jpx and Ftx both showed the potential to activate X-inactivation in mice^42,43^. But the functions of their human orthologs have not been studied^37^. The mean prediction scores of the transcribed regions in JPX and FTX are 0.256 and 0.304, respectively. This suggested only moderate functional potential of these two human lncRNAs. However, both scores received a substantial boost when the entire gene was considered. On Figure 4A, several functional peaks could also be clearly observed in the untranscribed regions in JPX and FTX. These results might guide the detection of novel regulatory elements in human XIC.

Besides the mentioned lncRNA genes, the human XIC also contains 6 protein-coding genes, NAP1L2, CDX4, CHIC1, ZCCHC13, SLC16A2, and RLIM. It is notable that most of their exons clearly reside in the functional peaks in Figure 4A, showing the ability of GenoCanyon to capture the functional landscape of this genomic region. We calculated the mean prediction scores for all the RefSeq transcripts of these genes. In CDX4, CHIC1, and SLC16A2, all the transcript scores were substantially larger than the scores of untranscribed regions. Among the 6 protein-coding genes, Rlim (also referred to as Rnf12) produces the U3 ubiquitin ligase that acts in a dose-dependent manner on the initiation of X-inactivation^46^. The human RLIM gene has a high mean predicted score 0.973. Two of its RefSeq transcripts both got 0.930 as the mean score, which is also very high. In Figure 4A, the RLIM gene perfectly lies in an isolated functional plateau, which suggests its strong functional potential in human. It has been observed that the homologous pairing of two regions (Tsix/Xite and Xpr/Slc16a2) might have impacts on Xist upregulation^47,48^. In human XIC, the TSIX/XITE region has been truncated, but the region surrounding the SLC16A2 gene showed its functional potential in our prediction. The exons of SLC16A2 lie in two large separate functional peaks, suggesting the importance of the transcribed region as well as a large bulk of untranscribed region in SLC16A2. Whether these regions serve as the human XPR remains to be investigated. More interestingly, 8 pathogenic SNPs in SLC16A2 have been submitted to ClinVar^49^.

These variants were believed to be involved in Allan-Herndon-Dudley syndrome, showing that SLC16A2 has its crucial function in other processes as well. The other genes in Xic have not been related to X-inactivation yet. Our prediction suggested that the exons of NAP1L2, CDX4, and CHIC1 all showed different levels of functional potential, which is not surprising because of their protein-coding nature. The human XIC also contains several microRNA genes and one pseudogene MAP2K4P1. MAP2K4P1 did not get a high score, which was in agreement with its pseudogene status. The microRNA transcript might partially explain the large functional plateau near the 5’end of FTX.

XACT, a recently discovered lncRNA coating the active X chromosome in human pluripotent cells, has been shown to take part in X-inactivation initiation uniquely in human^37,38^. It lies in a 1.7 Mb large intergenic region between protein-coding genes AMOT and HTR2C. A shorter transcript T113.3 upstream of XACT was also identified. But its function has not been studied. We investigated this region using GenoCanyon. The AMOT gene and the HTR2C exons both showed substantial functional potential. A clear plateau of high scores could also be observed in the intergenic domain (Figure 4B). The mean prediction score for the entire intergenic region, XACT, and T113.3 were 0.148, 0.383, and 1.000, respectively. Although the mean predicted score for XACT was only moderate, it still confirmed the functional signal in such a lowered-scored intergenic domain. Also, our prediction suggested the importance of T113.3 and its surrounding region.

### Investigating the Ability of Classifying Variants

GenoCanyon was not designed as a variant classifier. However, enrichment in prediction score is still expected for the known pathogenic variants. We downloaded all the annotated variants from ClinVar in June 2014^49^. The subset of single nucleotide variants annotated as “Pathogenic”, “Likely Pathogenic”, or “Pathogenic/Likely Pathogenic” was treated as the positive set. Similarly, the subset of SNVs annotated as “Benign”, “Likely Benign”, or “Benign/Likely Benign” was treated as the negative set. The positive set contained 19,242 variants, and the negative set contained 8,874 variants. The mean prediction score in the positive set and the negative set were 0.912 and 0.735, respectively. When using 0.5 as the natural cut-off, the sensitivity was as high as 0.915, with a low specificity of 0.263. The AUC was 0.727.

It is worth noting that GenoCanyon measures the functional potential of genomic locations, not the tolerability of specific variants. The transcribed regions in a crucial protein-coding gene should be expected to have a high functional score. However, it would still be natural to observe many tolerable synonymous SNPs in that gene. All these tolerable SNPs become “false-positives” in the analysis above, leading to a low specificity. Moreover, many of the known “benign” variants are by-products of association studies. Their properties were investigated because they lie in candidate regions in the disease pathway, which explains why the mean prediction score of benign variants was also high. On the other hand, if a variant were shown to be pathogenic in experiments, the underlying region would surely have some functions related to the disease. In this sense, the high sensitivity of GenoCanyon suggests that it may be a good indicator of its prediction ability. Finally, the performance of supervised-learning-based methods is highly sensitive to the choice of training data. For example, when using common variants with matched regions as the negative training set, the performance of GWAVA on its own training data dropped substantially (AUC=0.71)^14^.

## Discussion

The HBB gene cluster, ZRS, and the X-inactivation center all have been paradigms for studying the complex genomic regulatory network. The prediction results in these regions showed that GenoCanyon is capable of detecting functional regions in the human genome, which is a unique feature most existing whole-genome annotation tools don’t have. With the wide adoption of next-generation sequencing, GenoCanyon may help researchers focus on candidate regions that are likely to be functional and reduce the spurious signals among the overwhelming genomic information.

Throughout this article, we have discussed the differences between GenoCanyon and variant classifiers in that GenoCanyon measures the functional potential of genomic locations instead of the pathogenicity of a specific variant and a high score does not necessarily imply deleteriousness. However, in some scenarios that variants distribute across the entire genome, GenoCanyon may still serve well as a conservative tool for noise reduction. For example, sequencing technology is rapidly becoming a focus of efforts in genomic epidemiology. However, the overwhelming number of rare variants in the human genome brings the issue of extreme multiple testing. It has been discussed recently that the sample size required for a well-powered RVAS (rare variants association study) using sequencing is similar to that of a traditional GWAS (genome-wide association study)^50^. Without a huge cohort, true signals could be easily overshadowed by extreme yet spurious observations. In this case, GenoCanyon could be used to filter the SNPs and reduce 2/3 of the tests as more than 2/3 of the human genome is less likely to be functional. Moreover, the high sensitivity of GenoCanyon ensures that the true signal is still kept in the dataset. The ability of predicting functional potential at each nucleotide is another useful feature of GenoCanyon. In association studies, genetic variants are used as markers capturing signals for nearby regions. Therefore, for each SNP, the mean prediction score for its surrounding region may serve well as a prior in post-GWAS prioritization. Existing variant classifiers cannot achieve this task because they only predict the deleteriousness of genotyped variants. It is worth noting that most of the annotation data have a resolution ranging from tens to hundreds of nucleotides due to the limitation of current experimental techniques. However, data input of these annotations is at nucleotide level, which makes it possible to measure the functional potential for each base pair.

Based on unsupervised learning, GenoCanyon does not suffer from the highly biased knowledge of the non-coding DNA. More importantly, the model can be generalized in many directions. Firstly, the ENCODE annotations used in GenoCanyon were clustered across several or even nearly a hundred different cell lines. Therefore, the current functional regions predicted by GenoCanyon are in fact the union of functional elements in different cell types. Using the annotations for one single cell type, a cell type-specific functional prediction tool could be built under the same framework. In studies where several candidate cell types are of interest, prediction based on the cell type-specific models would have higher specificity. Secondly, the model can be extended to other species. The functional elements in model organisms are generally better studied. Such tools for different species could potentially benefit the multi-species comparison and help detecting functional orthologs in human. Thirdly, in order to simplify the model, we transformed the biochemical annotations into binary variables (See **Methods**). Therefore, the information of signal strength has not been used. When these information as well as more annotations are incorporated using more complex modeling techniques, the specificity may be improved. Finally, the current model assumes the leading role of genetic function, and treats conservation measures and the biochemical signals as consequences. Among different annotations, conditional independence was also assumed (**Figure 1**). However, it would be interesting to investigate the correlations among variables in either the functional or the non-functional group. In that case, statistical graphical models could be implemented to make the model more flexible. These are all very interesting directions to generalize GenoCanyon. However, complex models lead to higher variance, intensive computation, and less interpretability. Dealing with these trade-offs has never been trivial. The good prediction results have shown that GenoCanyon has reached a nice balance. The current powerful features as well as its generalizable potential make GenoCanyon a unique and useful tool for whole-genome annotation.

## Methods

### Statistical Model

For each location in the human genome, define Z to be the latent indicator of function, where Z=1 indicates that location is functional and 0 otherwise. We selected 22 different annotations corresponding to either conservation score or biochemical activity, including 2 genomic conservation measures, 2 indicators of open chromatin, 8 histone modifications, and 10 TFBS peaks (Supplementary Table 1). These annotations are selected because their functional impacts are relatively well studied and easier to model. DNA methylation is not included in the model because the gene silencing mechanism requires modeling the functional impact of methylation to other nucleotides that are possibly far away, which is a challenging task. Genomic data for all the 22 annotations were downloaded from the UCSC Genome Browser except GERP (Supplementary Table 3). We denote the vector of all the annotations as ***A***.

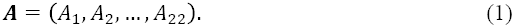

When a genomic location is functional (Z=1), we assume that the annotations have a joint probability density *f* (***A*|***Z* = 1); similarly, when a genomic location is non-functional (Z=0), we assume that the annotations have another joint density *f* (***A*|***Z* = 0). Since Z is unknown, the distribution of the observed data would be a mixture of *f* (***A*|***Z* = 1) and *f* (***A*|***Z* = 0). Instead of modeling direct causal relationships among these 22 annotations, we assume that they are connected only through Z. In other words, the 22 annotations are all modeled to be consequences of Z. Under these assumptions, the 22 different annotations are conditionally independent when Z is given^51^. Therefore, the conditional joint density of ***A*** given Z can be factorized as

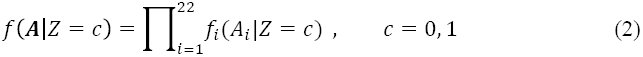

Finally, for a genomic location, assume π to be the prior probability of being functional, i.e.

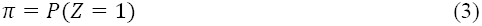

Then, given the annotations, the posterior probability of Z=1 can be used as a reasonable functional measure when the parameter estimates are plugged in.

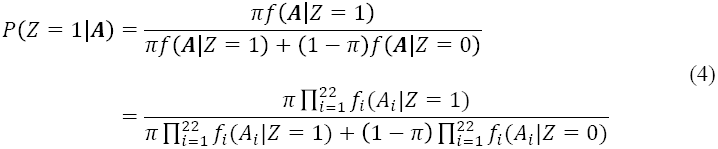

We chose GERP^52^ and PhyloP^53^ as the conservation measures because both of them are approximately normally distributed and therefore easier to model. PhyloP46way was chosen instead of PhyloP100way because a large phylogenetic distance would bring too little conserved signal as well as many incomplete data. All the other annotations were cell type-specific, so we coded them into binary variables to cluster the signal across cell lines. If signal was detected in at least one cell line, we coded the corresponding *A*_*i*_ = 1. Otherwise, *A*_*i*_ = 0. For DNase I, FAIRE, and TFBS, there were downloadable cluster files on the UCSC Genome Browser. A total of 125, 25, and 91 cell lines were clustered, respectively. We made our own histone peak cluster files across 16 cell lines from the Broad histone track on ENCODE (Supplementary Table 4). The 8 histone modifications were chosen because they are relatively well-studied^54^. We chose the top 10 Transcription Factors with the highest binding site coverage after being transformed into binary variables.

Finally, normal distribution and Bernoulli distribution were used to model the continuous and binary annotations, respectively.

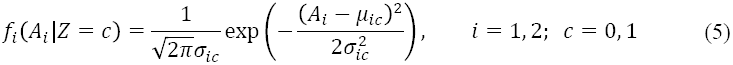

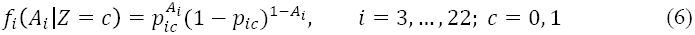

### Estimation

In total, our model has 49 parameters.

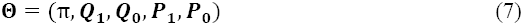

Where

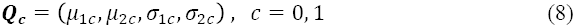

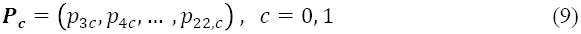

The GWAS Catalog^55^ was downloaded from the NHGRI GWAS Catalog website (http://www.genome.gov/gwastudies/) in July 2014. It contained 13,070 unique SNPs that were significant in GWAS studies. For each SNP, we marked the interval between its 500 bp upstream and 499 bp downstream. In this way, 13,070 intervals were collected. Each interval spanned 1k bp. After deleting the overlapping coordinates, the entire region spanned 12,801,840 bp. Each significant SNP in the GWAS Catalog hints the existence of functional elements nearby. These functional elements differ in their sizes and in the distance to the probed SNP. Since each interval was 1,000 bp in length and a large number of intervals were collected, the whole collection was a large enough and reasonably chosen set on which we could learn the distributions of annotations in both functional and non-functional groups. All the 22 annotations were then collected at each location in this set. The PhyloP scores and GERP scores were not available at 221,643 and 28,741 locations, respectively. After removing these locations, the final dataset contained 12,580,197 genomic locations. None of the other annotations have the issue of incomplete data. Finally, the Expectation-Maximization (EM) algorithm was used to estimate the parameters. As expected, the estimates showed solid differences between the functional and non-functional groups (Supplementary Table 5). We also tried replacing the missing conservation measures with the neutral score 0. Then the entire 12,801,840 locations were used to estimate the parameters. Little differences in parameter estimates were observed between the two approaches (Supplementary Table 6). Moreover, in order to test if the estimates are stable under different choices of datasets, we randomly sampled two subsets on chromosome 1, containing 2,000,000 and 6,000,000 bp, respectively. After adding these locations into the original 12,801,840 bp dataset, the parameters were estimated using the EM algorithm again. No substantial differences were observed in the estimates (Supplementary Tables 7 **and** 8). Based on these results, the GWAS-loci-based dataset containing 12,801,840 bp seems to contain enough functional elements for accurate parameter estimation, and is general enough so that genome heterogeneity does not have a strong impact on estimation. Finally, in order to check the sensitivity of our model to the perturbation in annotation data, we re-fitted the model multiple times after removing several annotations (Supplementary Table 9). The parameter estimates remained consistently stable in all these cases, suggesting that the framework we propose is robust to the choice of annotations. The stable estimates of marginal parameters also show that the potential correlations among annotations do not have a strong impact on model fitting.

### Marginal Effect of Different Annotations

For each binary annotation *A*_*j*_(j = 3, …, 22), its effect on the final prediction can be measured using the odds ratio.

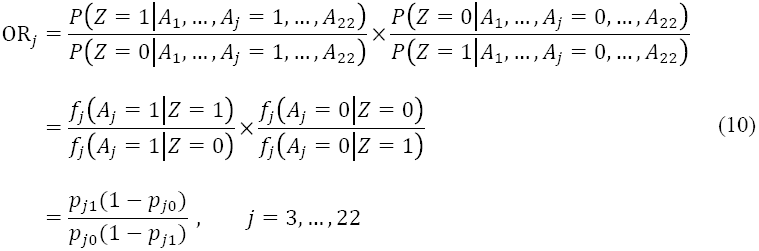

We calculated the odds ratios for all 20 binary annotations (Supplementary Table 5). The annotation with the least effect was the histone modification H3K27me3. According to our estimation, the probabilities to detect the H3K27me3 signal in functional and non-functional classes are almost the same (0.80 and 0.72). In fact, H3K27me3 has been discovered to be associated with Polycomb-repressed regions^56,57^, which could partially explain the phenomenon. All the other binary annotations showed variable yet substantial signals of function. The marginal effect of a continuous annotation depends on its value. The interpretation is also less straightforward. More importantly, although these statistics could help us gain some intuition of how each annotation works marginally, the final prediction relies on all of them. The effectiveness of the method needs to be tested as a whole.

In order to visualize the relative contribution of different sources of information (Figure 2C), posterior probabilities given a particular group of annotations were calculated for each location.

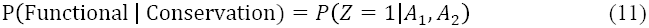

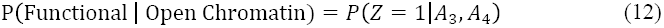

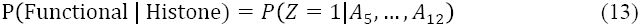

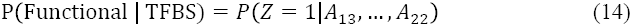

Then, for each CRM, the mean posterior probabilities were plotted.

### Estimating the Functional Proportion

After plugging in the parameter estimates, the prediction score could be calculated using formula (4). If the PhyloP or the GERP score was not available, the neutral value 0 was used. Using the cutoff 0.5, 33.3% of the human genome was predicted to be functional. However, it is notable that the EM algorithm also gave an estimate for the functional proportion, 42.7% in our case (Supplementary Table 5). This estimation was based on the 12,580,197 locations we chose, which might not represent the entire genome. 42.7% could be treated as the prior knowledge, but the final prediction will be driven by the actual annotations at each location. Therefore, 33.3% would still be a better estimation. To see if the prior had a strong effect, we estimated the functional proportion of chromosome 22 using different values for π while keeping other parameters unchanged. When using 0.3 and 0.5 as the π values, the estimated functional proportions were 0.376 and 0.389, respectively. Compared to the original estimate 0.383, there was not a substantial change.

### Figures and Web Application

All figures were plotted using R. The “ggbio” package was used to plot the chromosomes and transcripts^58^. The GenoCanyon web application was developed using the “shiny” package in R. The “bigmemory” package was implemented to access and manipulate massive datasets^59^. The GenoCanyon web application is available at http://genocanyon.med.yale.edu. The web server is implemented using Apache running on CentOS version 6.

## Acknowledgments

This study was supported by the National Institutes of Health grants R01 GM59507 and U01 HG005718, the VA Cooperative Studies Program of the Department of Veterans Affairs, Office of Research and Development, and the Yale World Scholars Program sponsored by the China Scholarship Council.

## Author Contributions

Q.L. and H.Z. designed the project. Q.L. wrote the initial draft and performed the analyses. Y.H. collected the annotation datasets. Q.L., J.S. and Y.C. developed the web server. K.C. advised on web server development. H.Z. advised on statistical and genetic issues.

## Supplementary Materials

**Supplementary Table 1.** The 22 annotations used in the model

| Notation | Annotation | Category |
| --- | --- | --- |
| $A_1$ | GERP | Conservation Measure |
| $A_2$ | PhyloP | |
| $A_3$ | DNase I | Open Chromatin |
| $A_4$ | FAIRE | |
| $A_5$ | H3k4me1 | Histone Modification |
| $A_6$ | H3k4me2 | |
| $A_7$ | H3k4me3 | |
| $A_8$ | H3k9ac | |
| $A_9$ | H3k27ac | |
| $A_{10}$ | H3k27me3 | |
| $A_{11}$ | H3k36me3 | |
| $A_{12}$ | H4k20me1 | |
| $A_{13}$ | CEBPB | TFBS |
| $A_{14}$ | CTCF | |
| $A_{15}$ | EP300 | |
| $A_{16}$ | FOS | |
| $A_{17}$ | GATA2 | |
| $A_{18}$ | JUND | |
| $A_{19}$ | MAX | |
| $A_{20}$ | MYC | |
| $A_{21}$ | POLR2A | |
| $A_{22}$ | RAD21 | |

**Supplementary Table 2.** Predicted functional proportion for each chromosome using 0.5 as the cutoff

| **Chromosome** | **Proportion** | **Chromosome** | **Proportion** |
| --- | --- | --- | --- |
| 1 | 0.332 | 13 | 0.365 |
| 2 | 0.331 | 14 | 0.344 |
| 3 | 0.356 | 15 | 0.392 |
| 4 | 0.293 | 16 | 0.338 |
| 5 | 0.374 | 17 | 0.334 |
| 6 | 0.316 | 18 | 0.362 |
| 7 | 0.321 | 19 | 0.313 |
| 8 | 0.328 | 20 | 0.340 |
| 9 | 0.388 | 21 | 0.331 |
| 10 | 0.337 | 22 | 0.383 |
| 11 | 0.322 | X | 0.281 |
| 12 | 0.307 | Y | 0.193 |
| **Overall** | **0.333** |  |  |

**Supplementary Table 3.**
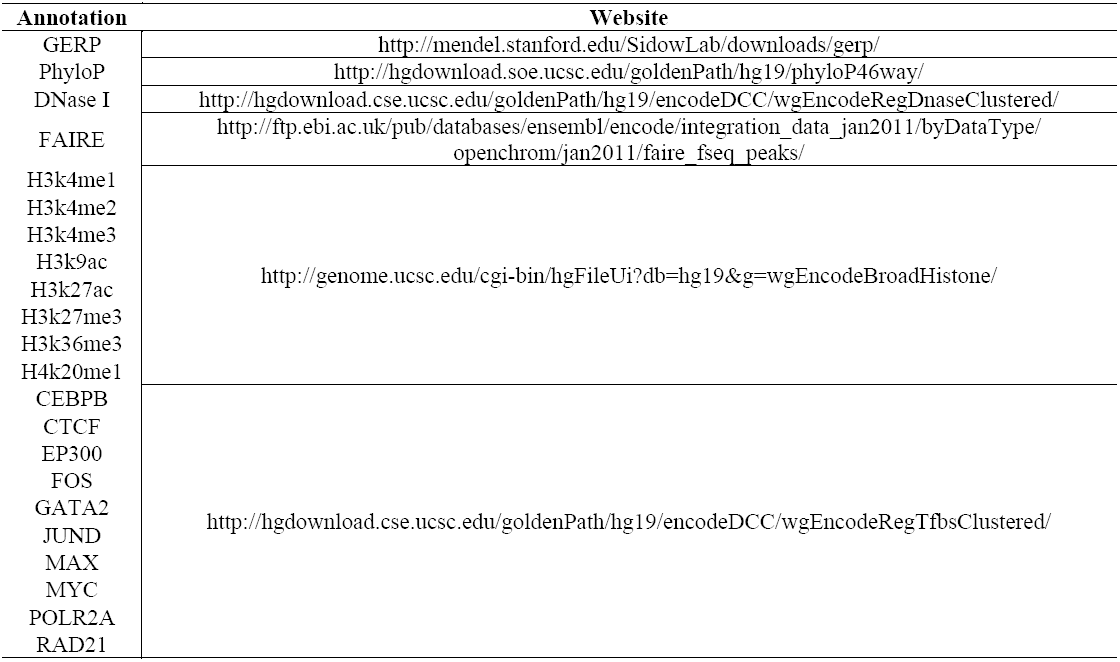
Online sources for the 22 annotations

**Supplementary Table 4.** 16 Cell lines used to cluster the histone peak signal

| Cell Line |
| --- |
| A549 |
| Dnd41 |
| Gm12878 |
| H1hesc |
| Helas3 |
| Hepg2 |
| Hmec |
| Hsmm |
| Hsmmt |
| Huvec |
| K562 |
| Monocd14ro1746 |
| Nha |
| Nhdfad |
| Nhek |
| Nhlf |

**Supplementary Table 5.** Estimates of all 49 parameters and the odds ratios for binary annotations

| Parameter | Estimate (Z=1) | Parameter | Estimate (Z=0) | Odds Ratio |
| --- | --- | --- | --- | --- |
| $\pi$ | 0.4274 | $1 - \pi$ | 0.5726 | - |
| $\mu_{11}$ | -0.1414 | $\mu_{10}$ | -0.0608 | - |
| $\mu_{21}$ | 0.1249 | $\mu_{20}$ | 0.0742 | - |
| $\sigma_{11}$ | 2.5274 | $\sigma_{10}$ | 1.6262 | - |
| $\sigma_{21}$ | 1.0765 | $\sigma_{20}$ | 0.6623 | - |
| $p_{31}$ | 0.3145 | $p_{30}$ | 0.0610 | 7.06 |
| $p_{41}$ | 0.3061 | $p_{40}$ | 0.0941 | 4.25 |
| $p_{51}$ | 0.9535 | $p_{50}$ | 0.3016 | 47.48 |
| $p_{61}$ | 0.7633 | $p_{60}$ | 0.0943 | 30.97 |
| $p_{71}$ | 0.6562 | $p_{70}$ | 0.0634 | 28.20 |
| $p_{81}$ | 0.7994 | $p_{80}$ | 0.1418 | 24.12 |
| $p_{91}$ | 0.8712 | $p_{90}$ | 0.2085 | 25.68 |
| $p_{10,1}$ | 0.7983 | $p_{10,0}$ | 0.7248 | 1.50 |
| $p_{11,1}$ | 0.8287 | $p_{11,0}$ | 0.3764 | 8.01 |
| $p_{12,1}$ | 0.9246 | $p_{12,0}$ | 0.6360 | 7.02 |
| $p_{13,1}$ | 0.0360 | $p_{13,0}$ | 0.0041 | 9.07 |
| $p_{14,1}$ | 0.0509 | $p_{14,0}$ | 0.0060 | 8.88 |
| $p_{15,1}$ | 0.0480 | $p_{15,0}$ | 0.0017 | 29.61 |
| $p_{16,1}$ | 0.0359 | $p_{16,0}$ | 0.0018 | 20.65 |
| $p_{17,1}$ | 0.0276 | $p_{17,0}$ | 0.0026 | 10.89 |
| $p_{18,1}$ | 0.0312 | $p_{18,0}$ | 0.0015 | 21.44 |
| $p_{19,1}$ | 0.0371 | $p_{19,0}$ | 0.0003 | 128.39 |
| $p_{20,1}$ | 0.0383 | $p_{20,0}$ | 0.0007 | 56.85 |
| $p_{21,1}$ | 0.1002 | $p_{21,0}$ | 0.0023 | 48.31 |
| $p_{22,1}$ | 0.0278 | $p_{22,0}$ | 0.0020 | 14.27 |

**Supplementary Table 6.** Estimates of all 49 parameters with the missing conservation values replaced by 0

| Parameter | Estimate (Z=1) | Parameter | Estimate (Z=0) |
| --- | --- | --- | --- |
| $\pi$ | 0.4305 | $1 - \pi$ | 0.5695 |
| $\mu_{11}$ | -0.1676 | $\mu_{10}$ | -0.0377 |
| $\mu_{21}$ | 0.1142 | $\mu_{20}$ | 0.0791 |
| $\sigma_{11}$ | 2.5549 | $\sigma_{10}$ | 1.5445 |
| $\sigma_{21}$ | 1.0886 | $\sigma_{20}$ | 0.6269 |
| $p_{31}$ | 0.3051 | $p_{30}$ | 0.0621 |
| $p_{41}$ | 0.2976 | $p_{40}$ | 0.0949 |
| $p_{51}$ | 0.9493 | $p_{50}$ | 0.3646 |
| $p_{61}$ | 0.7789 | $p_{60}$ | 0.1233 |
| $p_{71}$ | 0.6659 | $p_{70}$ | 0.0765 |
| $p_{81}$ | 0.8024 | $p_{80}$ | 0.1688 |
| $p_{91}$ | 0.8552 | $p_{90}$ | 0.2286 |
| $p_{10,1}$ | 0.8442 | $p_{10,0}$ | 0.7696 |
| $p_{11,1}$ | 0.8493 | $p_{11,0}$ | 0.4570 |
| $p_{12,1}$ | 0.9397 | $p_{12,0}$ | 0.7115 |
| $p_{13,1}$ | 0.0355 | $p_{13,0}$ | 0.0038 |
| $p_{14,1}$ | 0.0500 | $p_{14,0}$ | 0.0057 |
| $p_{15,1}$ | 0.0468 | $p_{15,0}$ | 0.0018 |
| $p_{16,1}$ | 0.0348 | $p_{16,0}$ | 0.0019 |
| $p_{17,1}$ | 0.0272 | $p_{17,0}$ | 0.0024 |
| $p_{18,1}$ | 0.0305 | $p_{18,0}$ | 0.0015 |
| $p_{19,1}$ | 0.0362 | $p_{19,0}$ | 0.0004 |
| $p_{20,1}$ | 0.0377 | $p_{20,0}$ | 0.0005 |
| $p_{21,1}$ | 0.0974 | $p_{21,0}$ | 0.0028 |
| $p_{22,1}$ | 0.0273 | $p_{22,0}$ | 0.0019 |

**Supplementary Table 7.** Estimates of all 49 parameters when an extra sample of randomly chosen 2,000,000 positions on chromosome 1 was added into the original dataset

| Parameter | Estimate (Z=1) | Parameter | Estimate (Z=0) |
| --- | --- | --- | --- |
| $\pi$ | 0.4332 | $1 - \pi$ | 0.5668 |
| $\mu_{11}$ | -0.1648 | $\mu_{10}$ | -0.0334 |
| $\mu_{21}$ | 0.1162 | $\mu_{20}$ | 0.0805 |
| $\sigma_{11}$ | 2.5571 | $\sigma_{10}$ | 1.5362 |
| $\sigma_{21}$ | 1.0876 | $\sigma_{20}$ | 0.6247 |
| $p_{31}$ | 0.3058 | $p_{30}$ | 0.0593 |
| $p_{41}$ | 0.2937 | $p_{40}$ | 0.0935 |
| $p_{51}$ | 0.9455 | $p_{50}$ | 0.3382 |
| $p_{61}$ | 0.7673 | $p_{60}$ | 0.1094 |
| $p_{71}$ | 0.6456 | $p_{70}$ | 0.0642 |
| $p_{81}$ | 0.7880 | $p_{80}$ | 0.1476 |
| $p_{91}$ | 0.8516 | $p_{90}$ | 0.2276 |
| $p_{10,1}$ | 0.8388 | $p_{10,0}$ | 0.7393 |
| $p_{11,1}$ | 0.8371 | $p_{11,0}$ | 0.4247 |
| $p_{12,1}$ | 0.9392 | $p_{12,0}$ | 0.6928 |
| $p_{13,1}$ | 0.0353 | $p_{13,0}$ | 0.0037 |
| $p_{14,1}$ | 0.0497 | $p_{14,0}$ | 0.0057 |
| $p_{15,1}$ | 0.0464 | $p_{15,0}$ | 0.0017 |
| $p_{16,1}$ | 0.0351 | $p_{16,0}$ | 0.0019 |
| $p_{17,1}$ | 0.0276 | $p_{17,0}$ | 0.0024 |
| $p_{18,1}$ | 0.0307 | $p_{18,0}$ | 0.0014 |
| $p_{19,1}$ | 0.0359 | $p_{19,0}$ | 0.0004 |
| $p_{20,1}$ | 0.0379 | $p_{20,0}$ | 0.0005 |
| $p_{21,1}$ | 0.0957 | $p_{21,0}$ | 0.0025 |
| $p_{22,1}$ | 0.0270 | $p_{22,0}$ | 0.0018 |

**Supplementary Table 8.** Estimates of all 49 parameters when an extra sample of randomly chosen 6,000,000 positions on chromosome 1 was added into the original dataset

| Parameter | Estimate (Z=1) | Parameter | Estimate (Z=0) |
| --- | --- | --- | --- |
| $\pi$ | 0.4393 | $1 - \pi$ | 0.5607 |
| $\mu_{11}$ | -0.1747 | $\mu_{10}$ | -0.0126 |
| $\mu_{21}$ | 0.1135 | $\mu_{20}$ | 0.0824 |
| $\sigma_{11}$ | 2.5745 | $\sigma_{10}$ | 1.4384 |
| $\sigma_{21}$ | 1.0908 | $\sigma_{20}$ | 0.5882 |
| $p_{31}$ | 0.3015 | $p_{30}$ | 0.0523 |
| $p_{41}$ | 0.2838 | $p_{40}$ | 0.0886 |
| $p_{51}$ | 0.9317 | $p_{50}$ | 0.2969 |
| $p_{61}$ | 0.7388 | $p_{60}$ | 0.0884 |
| $p_{71}$ | 0.6056 | $p_{70}$ | 0.0477 |
| $p_{81}$ | 0.7566 | $p_{80}$ | 0.1150 |
| $p_{91}$ | 0.8387 | $p_{90}$ | 0.2214 |
| $p_{10,1}$ | 0.8298 | $p_{10,0}$ | 0.6918 |
| $p_{11,1}$ | 0.8120 | $p_{11,0}$ | 0.3750 |
| $p_{12,1}$ | 0.9362 | $p_{12,0}$ | 0.6575 |
| $p_{13,1}$ | 0.0342 | $p_{13,0}$ | 0.0034 |
| $p_{14,1}$ | 0.0482 | $p_{14,0}$ | 0.0051 |
| $p_{15,1}$ | 0.0448 | $p_{15,0}$ | 0.0015 |
| $p_{16,1}$ | 0.0347 | $p_{16,0}$ | 0.0018 |
| $p_{17,1}$ | 0.0277 | $p_{17,0}$ | 0.0021 |
| $p_{18,1}$ | 0.0303 | $p_{18,0}$ | 0.0012 |
| $p_{19,1}$ | 0.0346 | $p_{19,0}$ | 0.0003 |
| $p_{20,1}$ | 0.0373 | $p_{20,0}$ | 0.0004 |
| $p_{21,1}$ | 0.0911 | $p_{21,0}$ | 0.0019 |
| $p_{22,1}$ | 0.0259 | $p_{22,0}$ | 0.0016 |

**Supplementary Table 9.** Estimates of parameters when GERP, DNase I, H3k4me2 and H4k20me1, and CEBPB and MAX are dropped from the model, respectively

| Param. | Estimate<br>(Z=1)<br>Without<br>GERP | Estimate<br>(Z=1)<br>Without<br>DNaseI | Estimate<br>(Z=1)<br>Without<br>H3k4me2<br>H4k20me1 | Estimate<br>(Z=1)<br>Without<br>CEBPB<br>MAX | Param. | Estimate<br>(Z=0)<br>Without<br>GERP | Estimate<br>(Z=0)<br>Without<br>DNaseI | Estimate<br>(Z=0)<br>Without<br>H3k4me2<br>H4k20me1 | Estimate<br>(Z=0)<br>Without<br>CEBPB<br>MAX |
| --- | --- | --- | --- | --- | --- | --- | --- | --- | --- |
| $\pi$ | 0.4201 | 0.4361 | 0.4417 | 0.4375 | $1 - \pi$ | 0.5799 | 0.5639 | 0.5583 | 0.5625 |
| $\mu_{11}$ | - | -0.1671 | -0.2468 | -0.1696 | $\mu_{10}$ | - | -0.0368 | 0.0277 | -0.0344 |
| $\mu_{21}$ | 0.1362 | 0.1131 | 0.0902 | 0.1126 | $\mu_{20}$ | 0.0637 | 0.0796 | 0.0974 | 0.0799 |
| $\sigma_{11}$ | - | 2.5469 | 2.6781 | 2.5510 | $\sigma_{10}$ | - | 1.5412 | 1.3288 | 1.5324 |
| $\sigma_{21}$ | 1.0328 | 1.0855 | 1.1355 | 1.0861 | $\sigma_{20}$ | 0.7009 | 0.6249 | 0.5433 | 0.6225 |
| $p_{31}$ | 0.3069 | - | 0.3039 | 0.3015 | $p_{30}$ | 0.0651 | - | 0.0582 | 0.0618 |
| $p_{41}$ | 0.3001 | 0.2933 | 0.2961 | 0.2948 | $p_{40}$ | 0.0967 | 0.0962 | 0.0921 | 0.0946 |
| $p_{51}$ | 0.9653 | 0.9484 | 0.9175 | 0.9467 | $p_{50}$ | 0.3633 | 0.3594 | 0.3779 | 0.3593 |
| $p_{61}$ | 0.7970 | 0.7736 | - | 0.7724 | $p_{60}$ | 0.1219 | 0.1208 | - | 0.1202 |
| $p_{71}$ | 0.6817 | 0.6623 | 0.6425 | 0.6594 | $p_{70}$ | 0.0755 | 0.0733 | 0.0831 | 0.0742 |
| $p_{81}$ | 0.8181 | 0.7998 | 0.7802 | 0.7978 | $p_{80}$ | 0.1687 | 0.1645 | 0.1736 | 0.1645 |
| $p_{91}$ | 0.8725 | 0.8562 | 0.8357 | 0.8517 | $p_{90}$ | 0.2272 | 0.2215 | 0.2314 | 0.2235 |
| $p_{10,1}$ | 0.8438 | 0.8431 | 0.8423 | 0.8441 | $p_{10,0}$ | 0.7712 | 0.7697 | 0.7696 | 0.7688 |
| $p_{11,1}$ | 0.8574 | 0.8513 | 0.8390 | 0.8476 | $p_{11,0}$ | 0.4581 | 0.4515 | 0.4573 | 0.4534 |
| $p_{12,1}$ | 0.9417 | 0.9398 | - | 0.9394 | $p_{12,0}$ | 0.7141 | 0.7091 | - | 0.7089 |
| $p_{13,1}$ | 0.0361 | 0.0347 | 0.0352 | - | $p_{13,0}$ | 0.0039 | 0.0041 | 0.0034 | - |
| $p_{14,1}$ | 0.0505 | 0.0483 | 0.0501 | 0.0492 | $p_{14,0}$ | 0.0061 | 0.0066 | 0.0047 | 0.0058 |
| $p_{15,1}$ | 0.0477 | 0.0458 | 0.0460 | 0.0460 | $p_{15,0}$ | 0.0020 | 0.0021 | 0.0015 | 0.0018 |
| $p_{16,1}$ | 0.0355 | 0.0338 | 0.0346 | 0.0342 | $p_{16,0}$ | 0.0020 | 0.0024 | 0.0014 | 0.0020 |
| $p_{17,1}$ | 0.0274 | 0.0266 | 0.0272 | 0.0268 | $p_{17,0}$ | 0.0027 | 0.0026 | 0.0019 | 0.0023 |
| $p_{18,1}$ | 0.0311 | 0.0300 | 0.0302 | 0.0300 | $p_{18,0}$ | 0.0015 | 0.0016 | 0.0011 | 0.0015 |
| $p_{19,1}$ | 0.0370 | 0.0357 | 0.0355 | - | $p_{19,0}$ | 0.0004 | 0.0005 | 0.0003 | - |
| $p_{20,1}$ | 0.0385 | 0.0370 | 0.0369 | 0.0370 | $p_{20,0}$ | 0.0006 | 0.0006 | 0.0004 | 0.0006 |
| $p_{21,1}$ | 0.0994 | 0.0958 | 0.0958 | 0.0960 | $p_{21,0}$ | 0.0030 | 0.0031 | 0.0022 | 0.0028 |
| $p_{22,1}$ | 0.0276 | 0.0263 | 0.0275 | 0.0269 | $p_{22,0}$ | 0.0021 | 0.0024 | 0.0012 | 0.0019 |

**Supplementary Figure 1.**
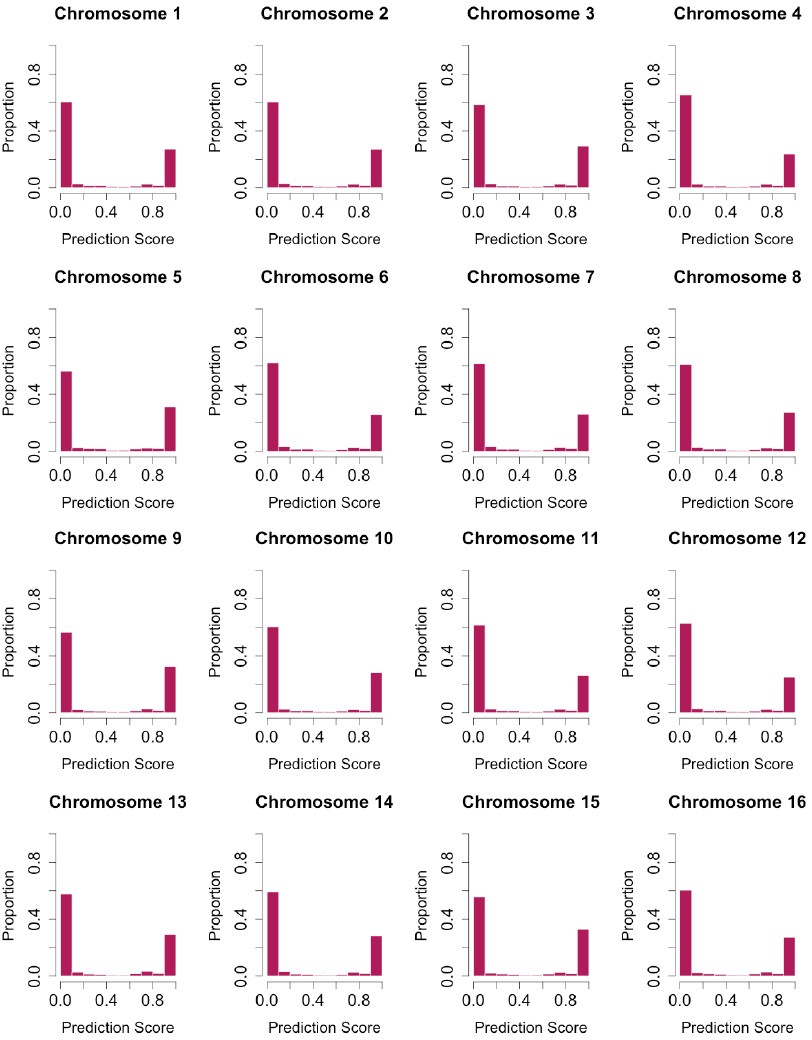
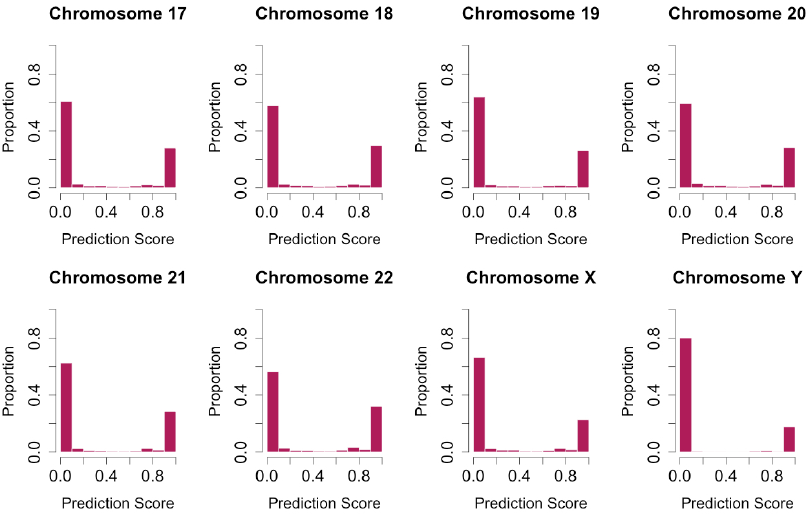
Histograms of prediction score in 24 chromosomes.

